# INGEsT: An Open-Source Behavioral Setup for Studying Self-motivated Ingestive Behavior and Learned Operant Behavior

**DOI:** 10.1101/2024.03.10.584229

**Authors:** Zhe Zhao, Binbin Xu, Cody L. Loomis, Skylar A. Anthony, Isaac McKie, Abhishikta Srigiriraju, McLean Bolton, Sarah A. Stern

## Abstract

Ingestive behavior is driven by negative internal hunger and thirst states, as well as by positive expected rewards. Although the neural substrates underlying feeding and drinking behaviors have been widely investigated, they have primarily been studied in isolation, even though eating can also trigger thirst, and vice versa. Thus, it is still unclear how the brain encodes body states, recalls the memory of food and water reward outcomes, generates feeding/drinking motivation, and triggers ingestive behavior. Here, we developed an INstrument for Gauging Eating and Thirst (INGEsT), a custom-made behavioral chamber which allows for precise measurement of both feeding and drinking by combining a FED3 food dispenser, lickometers for dispensing liquid, a camera for behavioral tracking, LED light for optogenetics, and calcium imaging miniscope. In addition, *in vivo* calcium imaging, optogenetics, and video recordings are well synchronized with animal behaviors, e.g., nose pokes, pellet retrieval, and water licking, by using a Bpod microprocessor and timestamping behavioral and imaging data. The INGEsT behavioral chamber enables many types of experiments, including free feeding/drinking, operant behavior to obtain food or water, and food/water choice behavior. Here, we tracked activity of insular cortex and mPFC Htr3a neurons using miniscopes and demonstrate that these neurons encode many aspects of ingestive behavior during operant learning and food/water choice and that their activity can be tuned by internal state. Overall, we have built a platform, consisting of both hardware and software, to precisely monitor innate ingestive, and learned operant, behaviors and to investigate the neural correlates of self-motivated and learned feeding/drinking behaviors.

## Introduction

Ingestive behavior is critical to maintain the body’s energy and fluid levels and is necessary for survival. Hunger and thirst, respectively, induce feeding and drinking behaviors by negative reinforcement (Allen et al., 2017; Betley et al., 2015), whereas the rewarding effects of food and drink promote ingestive behavior through positive reinforcement (Stern et al., 2020). In addition, memory also impacts food and water consumption based on previous experience of behavioral consequences (Stern et al., 2021; Stern *et al*., 2020). However, it is mostly unknown how the brain encodes food and water deprivation compared to memories related to feeding and drinking, and how this generates motivation for ingestion. Specifically, it is unclear whether different neural dynamics underlie 1) food- and water-deprivation states, 2) motivation to seek food and water, and 3) food and water rewards. Understanding these mechanisms underlying ingestive behavior will better help to develop approaches to treat obesity and eating disorders, e.g., anorexia nervosa and bulimia nervosa.

The brain senses body energy and fluid states by peripheral hormones and the nervous system (Gizowski and Bourque, 2018; Leib et al., 2016; Rowland, 2004; Watts et al., 2022). The hypothalamus and hindbrain are key brain regions that receive hormonal signals as well as information arriving from vagus nerve. Subsequently, internal body state information reaches cortical regions through the thalamus. The insular cortex (or insula) and medial prefrontal cortex (mPFC) are reported as key interoceptive brain areas to sense internal body states and generate proper behaviors (Craig, 2003; Livneh and Andermann, 2021; Zhao et al., 2022; Zhao et al., 2020). *In vivo* imaging in the insula and electrophysiological recording in the mPFC showed specific feeding-response and drinking-response neurons (Eiselt et al., 2021; Livneh et al., 2017; Livneh et al., 2020). Furthermore, insular neurons also represent body food- and water-deprivation states (Livneh *et al*., 2017; Livneh *et al*., 2020). However, in these studies, mice under food- or water-restricted states are tested on different days with only food or only water. Thus, the neural dynamics of state transitions (e.g., hunger to thirst after eating food and thirst to hunger after drinking water), are unknown. One recent study developed a food/water choice behavioral task, in which food- or water-deprived head-restrained mice chose food and water on the left or right side after an odor stimulus (Richman et al., 2023). Electrophysiological recording in multiple brain regions including prefrontal and motor cortices, forebrain, and midbrain simultaneously reveals that different neurons correlate with specific phases of the trial. Some neurons displayed persistent activity throughout trials, but the activity patterns were different in the food/water trials, suggesting that these neurons represent the internal need state of the body (Richman *et al*., 2023). However, this study has several caveats: first, it used liquid food with additional sodium to motivate drinking rather than standard chow. Secondly, food/water on the right or left side cannot exclude the potential effect of direction factor on the neural activity. Last, the choice is triggered by an external cue instead of self-paced motivated behavior.

To overcome some of these shortcomings, we report here a novel setup, INstrument for Gauging Eating and Thirst (INGEsT), a custom-made feeding/drinking behavioral box to investigate the above questions. Using this chamber, we can observe feeding and drinking behavior simultaneously in combination with *in vivo* calcium imaging, which will allow for more naturalistic behavior, from both the standpoint of intermingling need states, as well as allowing for freely moving behavior. In addition to describing the hardware, we also establish a platform for animal behavior and imaging data analysis.

## Results

### INGEsT behavioral chamber and affiliated setups

To study feeding and drinking behavior in the same context, we developed a behavioral chamber, consisting of a pellet dispenser (FED3) and a two-port lickometer, to precisely record time points of pellet retrieval and licking water (**Figure 1A**). We called this chamber INstrument for Gauging Eating and Thirst (INGEsT). The affiliated setups include Inscopix miniscope, a voltage pulse generator (Pulse Pal), and two cameras (**Figure 1B**). FED3 is an open-source device that is widely used to measure food intake hourly or daily for free-feeding studies(Matikainen-Ankney et al., 2021). In addition, it has two nose poke ports for progressive ratio and similar operant tasks to study motivation and learning. The lickometers can detect the signal of licking. The nose poke, pellet retrieval, and lick signals are recorded by a microprocessor, Bpod (**Figures 1A and C**), which can also send out TTL signals to trigger in vivo calcium imaging, LED light for optogenetic perturbation, and video recordings of animal behavior. Calcium imaging data is synchronized with behavioral data by timestamps. Pulse Pal can be triggered to start stimulation with the protocol (specific frequency and duration of stimulation) set in the device (**Figure 1B and C**). We show photos of the mouse working area, FED3 and lickometer in the INGEsT, and Pulse Pal, Bpod, imaging acquisition box, etc. outside of the INGEsT (**Figures 1D-F**).

**Figure 1.**
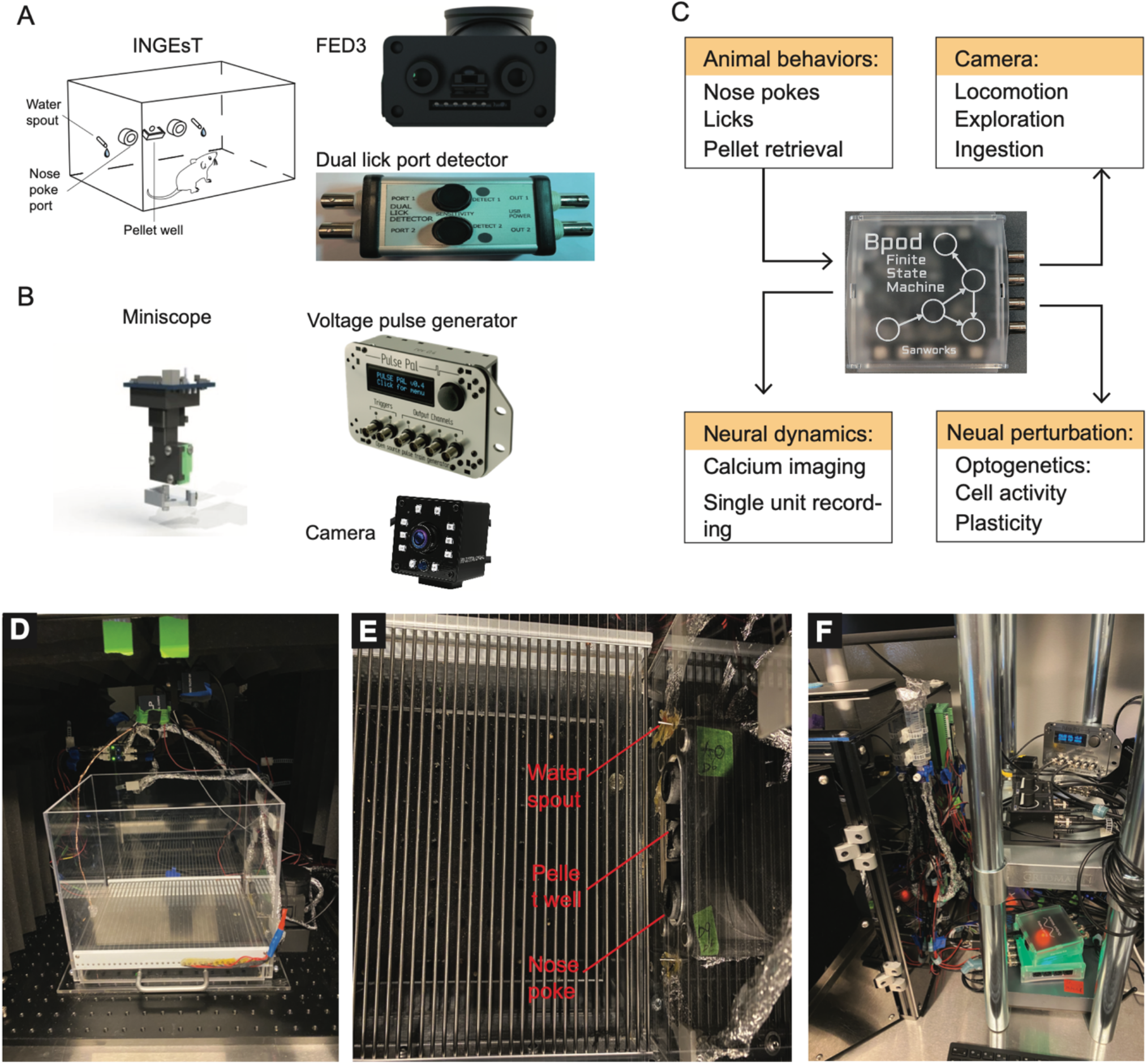
Overview of the INGEsT and other synchronized setups. **(A)** Main components of INGEsT: FED3 and a dual lick port detector. **(B)** Tools for studying neural activity and animal behavior: miniscope calcium imaging with freely moving mice, voltage pulse generator to trigger light for optogenetics, and a camera to record animal behavior. **(C)** Bpod microprocessor for integration and synchronization of INGEsT, miniscope, voltage pulse generator, and camera. **(D-F)** Pictures of real setup in our lab. **(D)** Animal working area in the INGEsT. **(E)** FED3 and water spout in the INGEsT. **(F)** Affiliated the setups, e.g., Bpod and Pulse Pal.

### Free feeding/drinking behavior and video recording

To validate that the setup can precisely record pellet retrieval and licking signals, we first measured feeding and drinking behaviors under food restriction (**Figure 2A**). Interestingly, on the first day, mice explored the new environment, and showed licking water spout before getting pellets even though they were food deprived (**Figures 2B-D**), suggesting that mice need to learn where to obtain food and water before they can appropriately satisfy their need states. After this first day, when mice were familiar with the behavioral chamber, food-deprived mice first ate pellets and then began to drank water only at the end of the training session. We found that mice tend to switch between feeding and drinking behaviors after 3 sessions of training (**Figures 2B and C**), about 10 to 20 minutes after starting the experiment (**Figure 2D**). Additionally, we recorded the animal behavior with a camera on the top of the INGEsT chamber (**Figures 2E-G**). Throughout the entire session, mice with miniscopes stayed primarily in the food and water area, whereas they spent little time in the center of the box (**Figures 2F-G**). We also analyzed the locomotion of mice without implants from the first day in the chamber. The mice increased the time in the food and water area after several days of experiments (**Figures 2H and I**). Thus, although the addition of miniscopes may lead to decreased movement in the chamber, animals are still able and willing to perform goal-directed behaviors, and will prioritize remaining in that zone.

**Figure 2.**
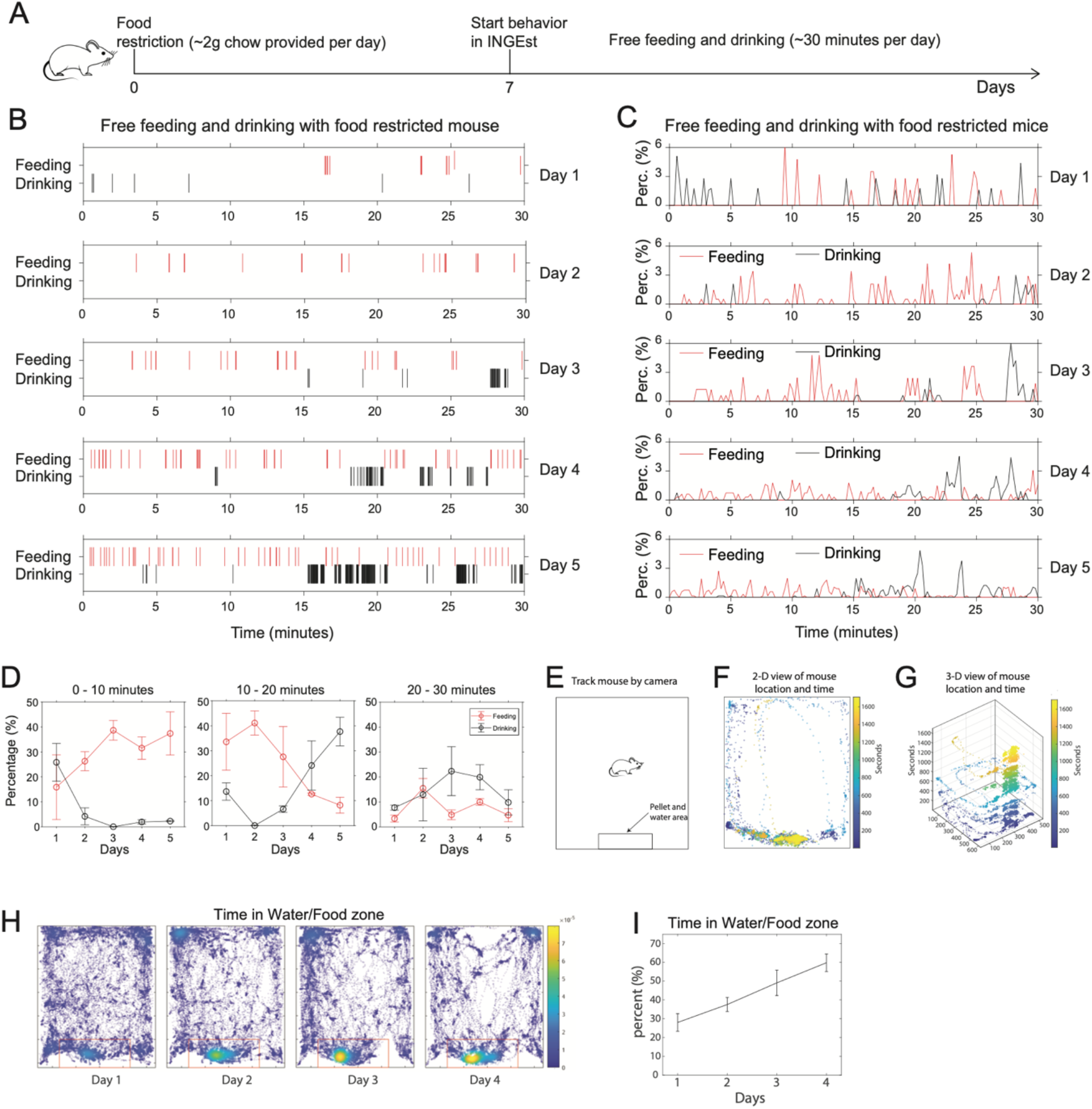
Free feeding and drinking behavioral patterns of food-restricted mice. **(A)** Scheme of free feeding and drinking behavioral protocol under food restriction. **(B)** Free feeding and drinking behaviors across days of one food-restricted mouse. **(C)** Free feeding and drinking behaviors across days of food-restricted mice (n = 5). **(D)** Different periods (0 – 10, 10 – 20, 20 – 30 minutes) of feeding and drinking behaviors across days (mice n = 5). **(E-G)** Animal location and time in INGEsT. **(E)** Scheme of the behavioral area in INGEsT. **(F)** 2-D view of the animal location and time in the INGEsT. **(G)** 3-D view of the animal location over time in the INGEsT. **(H)** Animal locomotion in the INGEsT across days. **(I)** Time in the water/food zone (mice n = 3).

### INGEsT facilitates imaging of neural activity during freely moving reversal learning task associated with chow pellets or water

To verify the synchronization of animal behavior with imaging data, we examined nose-poke associated pellet delivery in combination with in vivo calcium imaging with the Inscopix miniscope. We expressed a calcium indicator, GCaMP8m, in the insular cortex by using a viral approach (pGP-AAV-syn-jGCaMP8m). Immediately following the viral injection, we implanted a GRIN lens 50 μm above the injection site (**Figure 3A**). 6 weeks later after the surgery, we tested the calcium signal and started nose-poke associated pellet training (**Figures 3B and C**). This nose-poke training lasted for 30 minutes per day. The mice learned to make nose pokes for food on the first day of training and were able to perform at least 15 trials in 30 minutes on the following days, at which point we considered them to have learned the behavior. To avoid habitual nose-poke behavior, we use a reversal operant behavioral task in which mice can obtain a pellet after an active nose poke on one side of the ports and can have 5 active nose pokes on the same side of the nose-poke port. After 5 active nose pokes, the nose-poke port becomes inactive and the other port switches to active from inactive (**Figure 3D**). This reversal learning training lasted for 30 minutes per day. After several days of training, mice increased the correct rate and trial number of reversals (**Figure 3E**).

**Figure 3.**
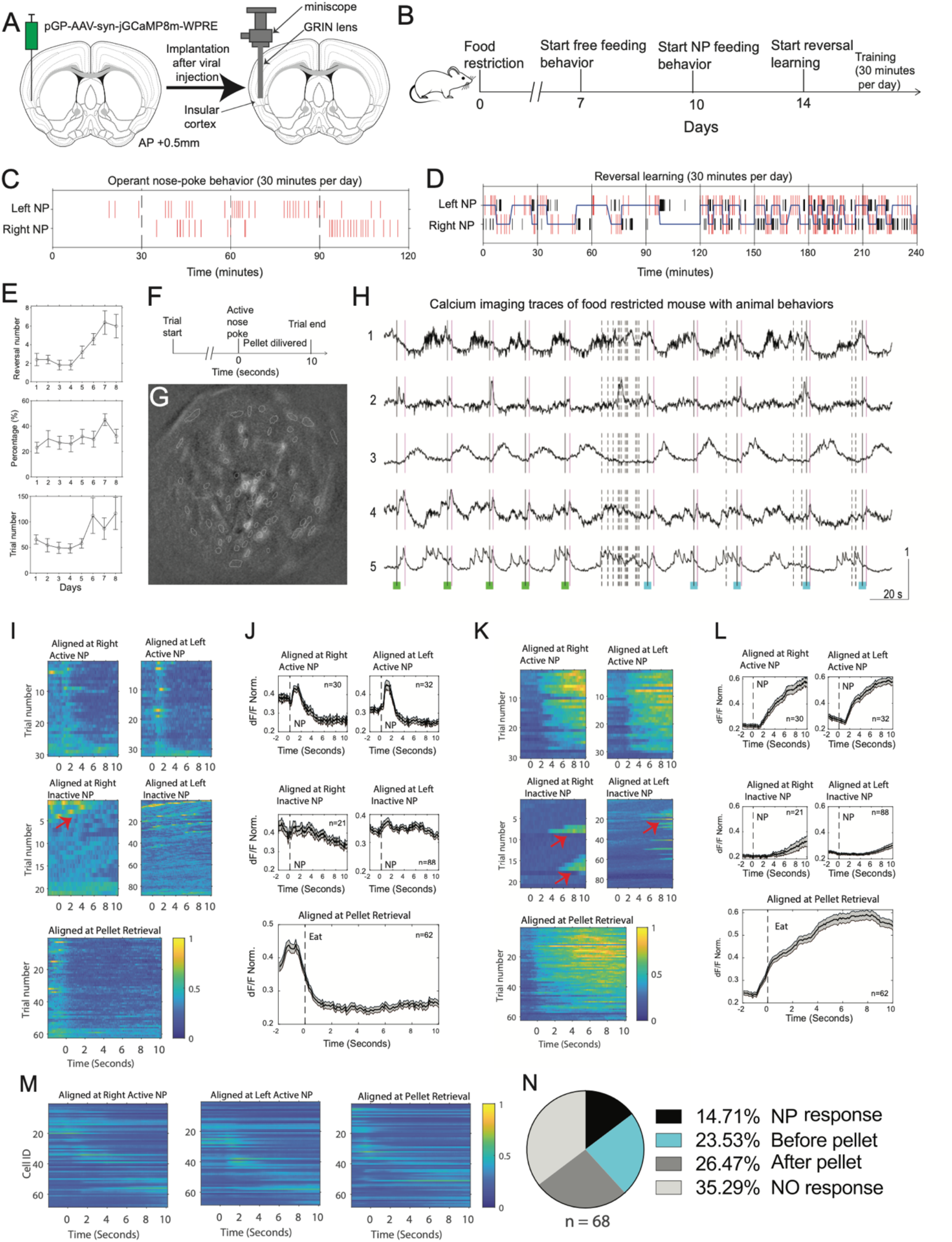
Reversal learning task associated with pellets and calcium imaging during the behavioral task. **(A)** Scheme of virus injection and GRIN lens implantation in the insular cortex. **(B)** Scheme of reversal learning behavioral training protocol with food-restricted mice. **(C)** Operant nose-poke behavior associated with food pellets under food restriction. Dashed lines represent the end of a daily training session. Red lines represent nose pokes on either the left or right side. **(D)** Reversal learning behavioral training with food pellets under food restriction. Nose-poke ports switch between active and inactive after 5 active nose pokes on the same side. The training lasts for 30 minutes per day, and requires at least 8 days. The blue line indicates which nose port is active, red lines indicate active nose pokes, black lines indicate inactive nose pokes. **(E-G)** Reversal learning across days. **(E)** Reversal number, Correct rate of nose pokes, Performed trial number (n=4). **(F)** Scheme of a trial, that lasts for 10 seconds after a nose poke at the active nose-poke port, in operant nose-poke training and reversal learning task. **(G)** Image of endoscope imaging view. **(H)** 5 traces of imaging coupled with animal behavior. Black lines: active nose pokes; magenta lines: licking water; Black dashed lines: inactive nose pokes; green square: right side of nose poke; cyan square: left side of nose poke. **(I-M)** Calcium imaging in the insular cortex during reversal operant behavior. Traces are aligned at time 0. **(I-J)** One example neuron that responds before pellet retrieval. **(K-L)** One example neuron that responds during pellet consumption. **(M)** The activity of all cells during specific types of trials. **(N)** Proportion of cells in response to a specific period in a trial.

In order to analyze the cell activity during different behaviors, e.g., nose poke, pellet retrieval, and pellet consumption, each active nose poke was set to make the nose-poke port inactive for the next 10 seconds (**Figure 3F**). We imaged 68 cells in the insular cortex during reversal learning training (**Figure 3G**), and we show 5 traces of neural activity from 5 representative cells during animal behavior (**Figure 3H**). The black and magenta lines on the trace indicate active nose pokes and pellet retrievals, respectively. The dashed lines indicate inactive nose pokes. In addition, the green and cyan squares indicate nose poke on the left side and right side of the nose-poke ports, respectively. Traces 1 and 5 show examples of neurons that increased activity before nose pokes, whereas traces 2 and 4 showed peak activity before pellet retrieval, and trace 3 showed an increase in activity after pellet retrieval (**Figure 3H**), suggesting different neurons encode different periods of the behavior. In other words, cells that are tuned to nose-poke, pellet retrieval, and pellet consumption after the retrieval are all present in the insular cortex. Furthermore, the behavioral data showed that the mouse made several inactive nose pokes before switching to the active nose-poke port (**Figure 3H**), which avoided habitual nose-poke behavior and also presented a reversal paradigm to investigate behavioral flexibility. To observe the neural response during specific periods of behavior, we analyzed neural activity 2 seconds before and 10 seconds after active/inactive nose pokes and pellet retrieval, and we show two examples of neural responses here (**Figures 3I-L**). Overall, the mouse performed 30 right and 32 left active nose pokes. However, it performed 21 right and 88 left inactive nose pokes, suggesting that the mouse exhibited a preference for left nose pokes. Accordingly, one neuron showed a peak of activity before pellet retrieval (**Figures 3I and J**), but the response was stronger in left nose-poke trials than in right nose-poke trials (**Figure 3J**). The arrows indicate the time points around the active nose pokes since the mouse can continue to do nose pokes (**Figure 3I**). Another cell showed a strong response during pellet consumption (**Figures 3K and L**). In this case, both left and right nose-poke trials showed a strong response during pellet consumption (**Figure 3L**). The arrows indicate the time points around the active nose pokes (**Figure 3K**) that are not themselves time-locked to inactive poking. To observe the different populations of neurons’ responses during the pellet associated nose-poke behavior, we made a heatmap of all cells with the same types of trials, e.g., left or right nose poke followed by pellet retrieval (**Figure 3M**). The results showed that different populations of neurons showed peak activity at different periods of the behavior. There were 14.71 percent of cells responding to nose pokes, 23.53 percent and 26.47 percent of cells responded to pellet retrievals and pellet consumption respectively; and 35.29 percent of cells were not tuned by the behavior (**Figure 3N**).

**I**NGEsT may also be used to study how the brain encodes operant drinking behavior. To demonstrate this, we next examined nose-poke associated water delivery in combination with freely moving *in vivo* calcium imaging (**Figure 4A**). Similarly to food-associated training, this nose-poke associated water training lasted for 30 minutes per day (**Figure 4B**). Here, the mice learned to complete nose pokes for water and perform at least 15 trials in a session on the first day of training, presumably because the mice have already completed the pellet-associated nose-poke behavior and are more familiar with the task and setup. Again, to avoid habitual nose-poke behavior, we use a reversal operant behavioral task which is similar to the pellet-associated reversal learning experiment. Mice obtain a drop of water if they lick the same side of the water spout with the nose-poke port after an active nose poke on one side of the port, and can have 5 active nose pokes on the same side of the nose-poke port. After 5 active nose pokes, the nose-poke port becomes inactive and the other port switches to active from inactive (**Figure 4C**). This reversal learning training lasted for 30 minutes per day. After several days of training, mice increased the correct rate and trial number of the reversals (**Figure 4D**).

**Figure 4.**
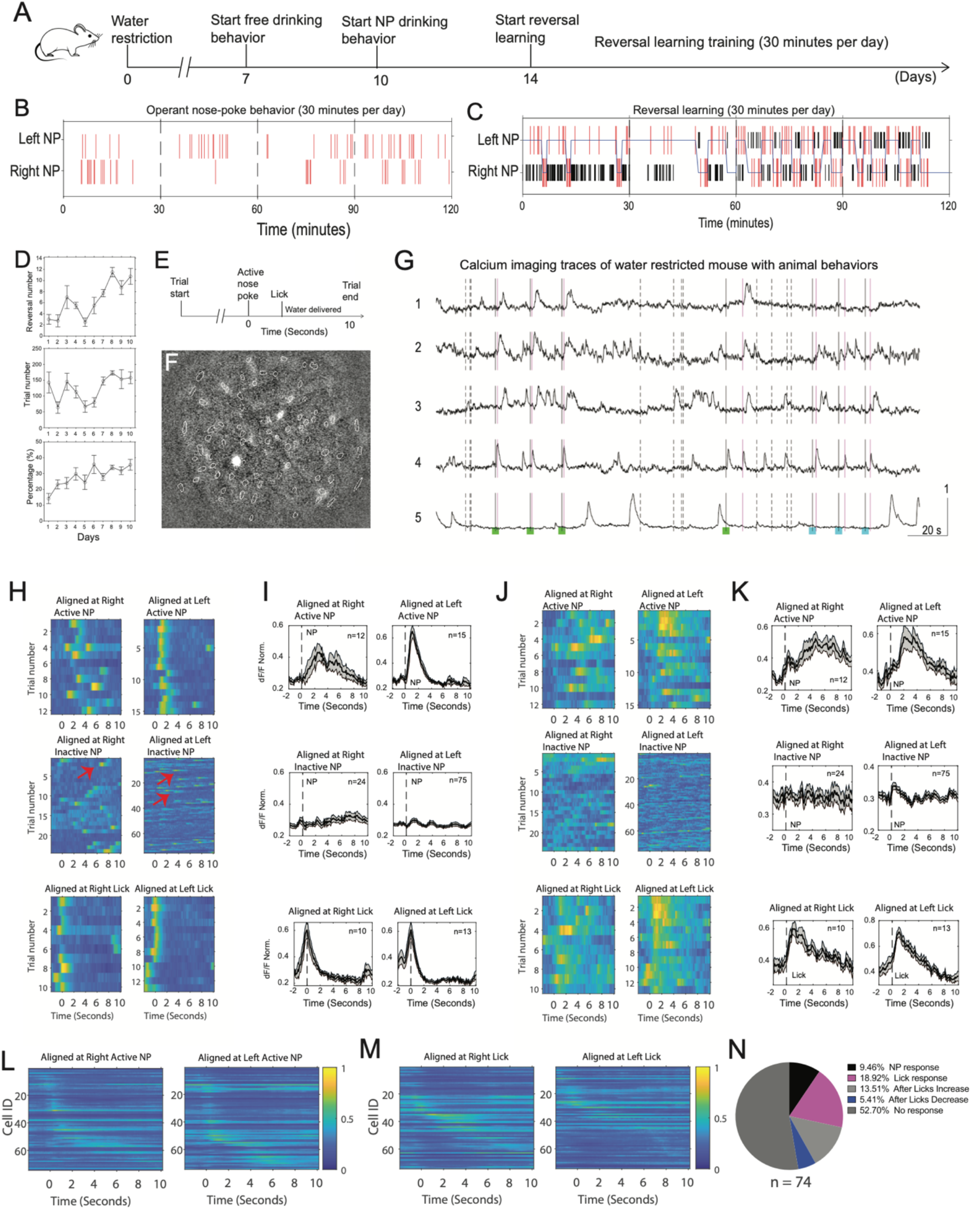
Reversal learning task associated with water and calcium imaging during the behavioral task. **(A)** Scheme of reversal learning behavioral training protocol with water-restricted mice. **(B)** Operant nose-poke behavior associated with water under water restriction. Red lines represent nose pokes on either the left or right side. **(C)** Reversal learning behavioral training with water under water restriction. Nose-poke ports switch between active and inactive after 5 active nose pokes on the same side. The training lasts for 30 minutes per day, and requires at least 8 days. The blue line indicates which nose port is active, red lines indicate active nose pokes, black lines indicate inactive nose pokes. **(D-G)** Reversal learning across days. **(D)** Reversal number, Correct rate of nose pokes, Performed trial number (n=4). **(E)** Scheme of a trial, that lasts for 10 seconds after a nose poke at the active nose-poke port, in operant nose-poke training and reversal learning task. **(F)** Image of endoscope imaging view. **(G)** 5 traces of imaging coupled with animal behavior. Black lines: active nose pokes; magenta lines: pellet retrieval; Black dashed lines: inactive nose pokes; green square: right side of nose poke; cyan square: left side of nose poke. **(H-K)** Calcium imaging in the insular cortex during reversal operant behavior. Traces are aligned at time 0. **(H-I)** One example neuron that responds to licking water. **(J-K)** One example neuron that responds during pellet consumption. **(L-M)** The activity of all cells during specific types of trials. **(N)** Proportion of cells in response to a specific period in a trial.

We then imaged the neural activity in the insular cortex during water reversal learning with the same parameters as for food (**Figure 4E**) in the same mouse as imaged in the food-associated reversal learning phase. Traces of 5 representative cells from 74 cells were shown with nose pokes and licks (**Figures 4F and G**). The black lines and magenta lines indicate active nose pokes and licks. The black dashed lines represent inactive nose pokes. In addition, the green and cyan squares indicate nose poke on left side and right side of nose-poke ports, respectively. Traces 1 and 3 showed neural responses after licks (or water rewards), whereas traces 2 and 4 showed peak activity at licks or water rewards. Trace 5 did not show a clear correlation with nose pokes and licks (**Figure 4G**), suggesting different neurons correlate with specific behaviors or behavioral outcomes. To observe neural responses in specific periods of the nose pokes followed by water reward, we aligned the imaging traces of calcium levels at nose pokes and licks (**Figures 4H and K**). This mouse performed 12 right and 15 left active nose pokes, but it performed 24 right and 75 left inactive nose pokes, again suggesting that this mouse exhibited a preference for the left nose poke port. Accordingly, one example neuron showed a peak of activity at the licking time (**Figures 4H and I**), but the response was stronger in left nose-poke trials than in the right nose-poke trials (**Figure 4I**). Another example neuron showed a strong response after water reward (**Figures 4J and K**). Both left and right nose-poke trials showed a strong response after licking water (**Figure 4K**). The peak of activity correlates with licking action since the licking lasts several seconds after the water reward. To observe activity of all imaged neurons in the same type of trials, we analyzed the calcium imaging traces of left and right active pokes. Most of imaged cells in the insular cortex showed peak activity around nose pokes and some cells showed responses around licks. Specifically, there were 9.46 percent of cells in response to nose pokes, 18.92 percent and 13.51 percent of cells showed peak activity at or after licks or water reward. Interestingly, 5.4 percent of cells showed a decrease in activity after licks or water reward. Therefore, these data show that this setup can be used to perform food- and water-associated operant behavior with *in vivo* calcium imaging to investigate neural mechanisms of drinking motivation, water reward, and reversal learning. Moreover, they indicate that although mice will nose-poke at the inactive port, the activity of some neurons indicates that the insular cortex can distinguish between nose pokes that will lead to reward and those that do not.

### INGEsT facilitates imaging of neural activity during freely moving feeding and drinking behavioral choice

To investigate if the same neurons encode different choices and rewards, we developed a self-motivated food and water choice behavior using INGEsT. These mice have completed the food/water-associated reversal learning task before the choice task (**Figure 5A**). In this task, both nose-poke ports are active, and active nose poke triggers pellet delivery and water availability. Both nose poke ports became inactive for 6 seconds after an active nose poke. Mice can then choose to retrieve a pellet or lick for water. If the mice retrieves the pellet, water will not be available until the next nose poke, but unretrieved pellets can remain in the food well even if mice choose to lick for water. We first trained the mice following long-term water restriction (0.8 ml of water is available daily). Following two days of training, (**Figure 5B**) the water-restricted mouse showed many water choice trials, but few food choice trials or water and food choice (defined as when the mouse chooses water, then food within 6 seconds after the previous nose poke), despite the fact that each nose poke triggers an available pellet. After 3 days of choice behavior, mice were switched from water restriction to food restriction (**Figures 5C and D**). Under food restriction condition, this mouse showed many food choice trials, but few water choice trials and few water and food choice trials (**Figure 5C**). As previously, this mouse exhibited a strong leftward bias. Interestingly, while water restricted, there were very few food-choice trials, but while food-restricted, there was an increase in the number of water+food choices (**Figure 5D**). This may reflect the mouse’s preference to wet the mouth for dry chow or to satisfy osmometric thirst as the mice eat more food.

**Figure 5.**
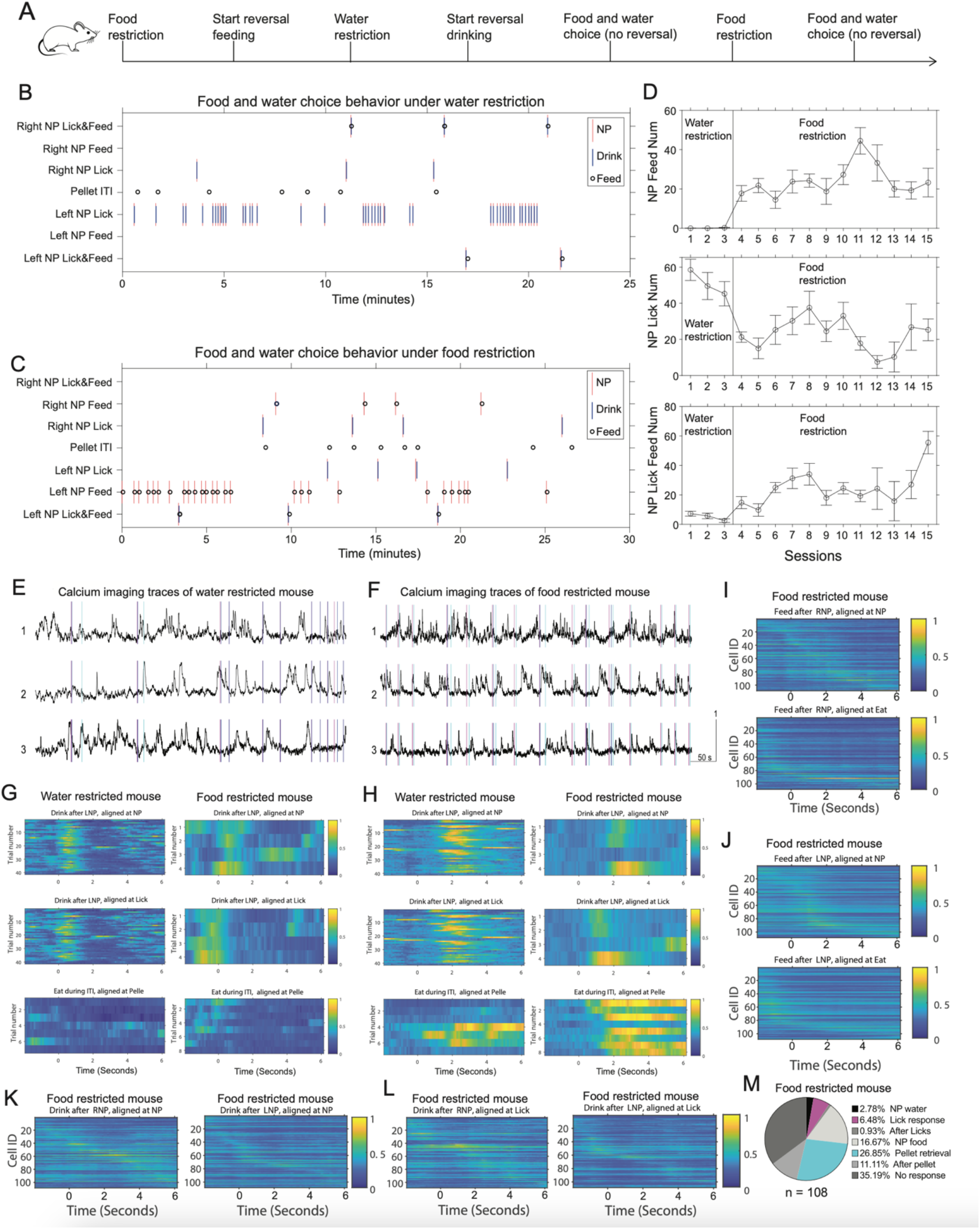
Feeding and drinking behavioral choice and calcium imaging during the behavioral task. **(A)** Scheme of reversal learning behavioral training protocol with water- or food-restricted mice. **(B)** Food and water choice behavior under water restriction. **(C)** Food and water choice behavior under food restriction. **(D)** Food and water choice behavior under water or food restriction across days (n=4). **(E-F)** 3 traces of imaging coupled with animal behavior from the same cells imaged on different days. Magenta lines: nose pokes; cyan lines: pellet retrieval; blue lines: licking water. **(G)** An example of a neuron that showed a peak of activity after the first lick of water and before pellet retrieval under the water restriction condition. This neuron showed a peak of activity in water choice trials before the first lick and before pellet retrieval. **(H)** An example of a neuron that showed a peak of activity after the first lick of water but not during the consumption of pellets. This neuron showed a higher response in water choice trials under the water restriction condition compared to the food restriction condition. This neuron showed a higher response in food choice trials under the food restriction condition compared to the water restriction condition. **(I-L)** The activity of all cells during specific types of trials. **(M)** Proportion of cells in response to a specific period in a trial.

Concurrent with choice behavior, we imaged neural activity in the insular cortex. We can detect feeding under water-restricted conditions and vice versa (**Figures 5E and F)**. In addition, we can compare the same cells imaged on different days. We showed activity traces of 3 cells under water restriction on the first day and under food restriction on the following day. Traces 1 of Figures 5 E and F are from the same cell as Figure 5G, and traces 2 of Figures 5 E and F are from the same cell as Figure 5H. We first compared the neural activity of the same cell in water choice trials and food choice trials, then compared the neural activity of the same cell under water restriction with food restriction (**Figures 5G and H**). One neuron showed a peak of activity after licking water but a weak response to pellet under water restriction conditions. During food restriction, this cell showed a peak of activity before licking water but no consistent response to pellet retrieval (**Figure 5G**), indicating this cell may represent specific water seeking behavior. Another cell showed peak activity after both licking water and pellet retrieval. The response during licking was stronger than pellet consumption under water restriction condition, and the response during licking was weaker than pellet consumption under food restriction conditions (**Figure 5H**), indicating this cell may represent general motivation. Furthermore, we analyzed neural activity of each cell in the same type of trials and showed the activity of all cells on a heatmap. Most cells showed peak activity around pellet retrieval (26.85%), and these cells are much more than other types of cells, e.g., nose poke responsive cells (16.67%), and lick-responsive cells (6.48%). (**Figures 5I-M**). The number of lick-responsive cells here (**Figure 5M**) are lower than during the imaging of water-restricted conditions (**Figure 4N**), indicating that the neural response is tuned by the internal body state.

### INGEsT facilitates imaging of neural activity in specific cell types during freely moving feeding and drinking behavioral choice

In the previous examples, we imaged from a general population of insular cortex neurons. To examine whether tuning of molecularly defined specific cell types can be detected in the choice task, we imaged one type of interneuron, 5-HT_3A_receptor-positive cells, in the medial prefrontal cortex (mPFC). We injected a viral vector carrying a Cre-dependent expression of GCaMP6s (pAAV.CAG.FLEX.GCaMP6s.WPRE.SV40) into the mPFC of a Htr3a-Cre:Ai-14 mouse, and the GRIN lens was implanted immediately after the virus injection (**Figure 6A**). This mouse completed the reversal nose-poke behavioral task with the same protocol as the previous experiments (**Figure 6B, and Figures 3-5**). We first performed feeding and drinking choice experiment for three continuous sessions under water restriction and then switched to food restriction (**Figure 6B**). Here, we showed the animal behavioral data on the third day of choice behavior under water restriction (**Figure 6C**) and started food restriction on the following day (**Figure 6C)**. We imaged 25 cells and 23 cells on the day of water restriction and food restriction, respectively. 3 calcium imaging traces with nose-poke behavior and food/water rewards under water restriction are shown (**Figure 6E**, top), and 3 traces from the same cells are shown to compare neural responses to the same behavior under different water- and food-restricted conditions (**Figure 6E**, bottom). The magenta lines indicate nose pokes. The blue and cyan lines represent licks and pellet retrievals, respectively. The neuron of trace 1 showed the peak activity is before licks under water restriction (**Figures 6F and G**), but not there under food restriction (**Figures 6H and I**), suggesting that the neural activity depends on internal body states. The neuron of trace 2 showed a strong response in the period of post-pellet retrieval, but a weak response to licking water (**Figures 6J and K**), indicating this neural response may also depend on the internal body state. To reveal the overview of neural responses of all imaged cells in specific types of trials, we averaged the imaging traces of the same types of trials from each cell, then made heatmap and sorted the cells based on the time of maximum value of the trace. It shows most of imaged cells showed peak activity around pellet retrievals under food restriction conditions (**Figures 6L and M**). There were 30.43 percent of cells responded to pellet retrieval, and 8.7 percent of cells showed a response at post-pellet retrieval.

**Figure 6.**
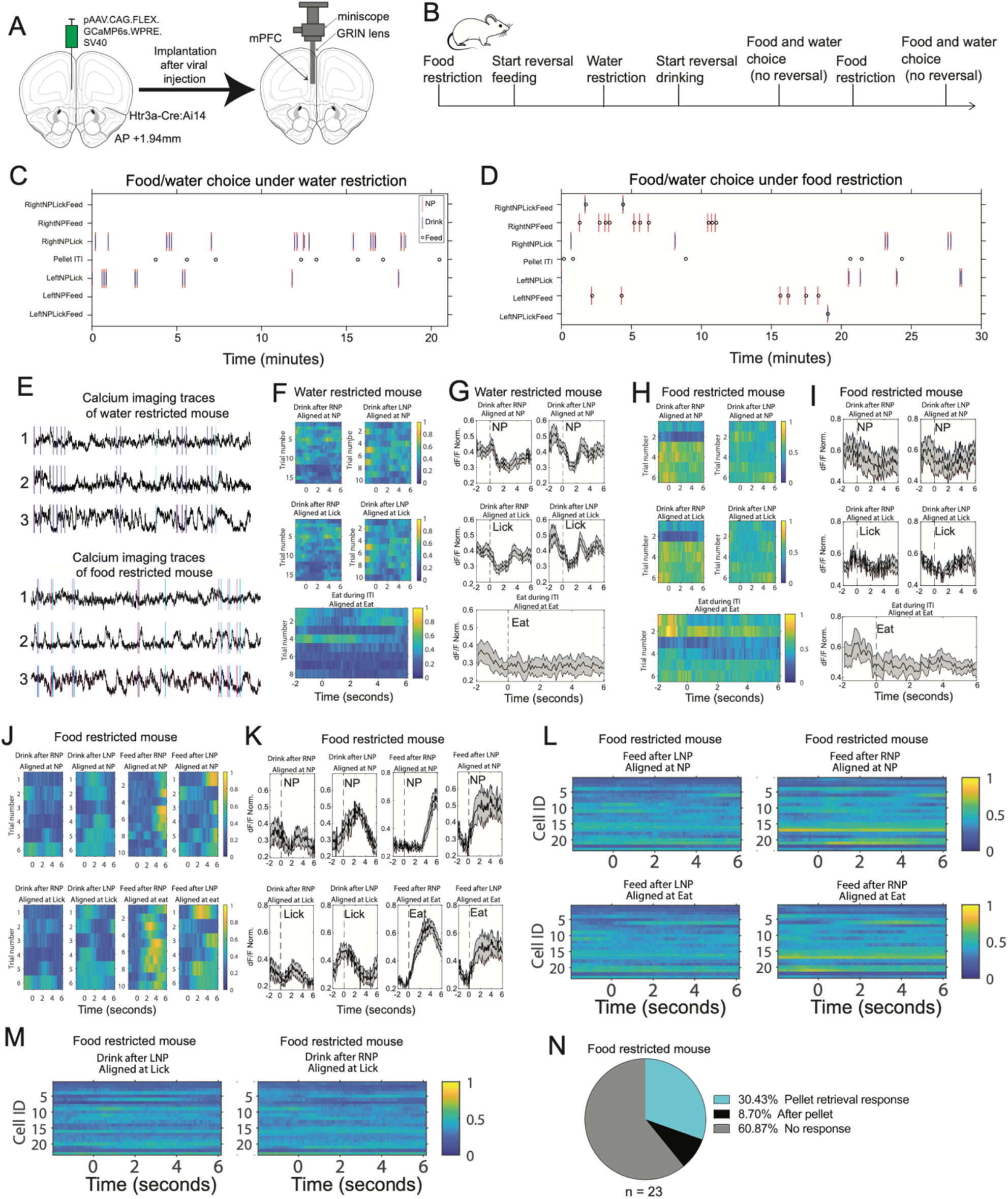
Feeding and drinking behavioral choice and calcium imaging of specific cell type in the medial prefrontal cortex during the behavioral task. **(A)** Scheme of virus injection and GRIN lens implantation in the insular cortex. **(B)** Scheme of reversal learning behavioral training protocol with water- or food-restricted mice. **(C)** Food and water choice behavior under water restriction. **(D)** Food and water choice behavior under food restriction. **(E)** 3 traces of imaging coupled with animal behavior from the same cells imaged on different days. Magenta lines: nose pokes; cyan lines: pellet retrieval; blue lines: licking water. **(F-I)** An example of a neuron that showed a decrease of activity after an active nose poke under the water restriction condition **(F-G)**, but did not show a change under food restriction conditions **(H-I). (J-K)** An example of a neuron that showed a peak of activity after the first lick of water and after pellet retrieval under the food restriction condition. The neural response after pellet retrieval is much stronger than the response after licking water. **(L-M)** The activity of all cells during specific types of trials under the food restriction condition. **(N)** Proportion of cells in response to a specific period in a trial.

## DISCUSSION

This study reports a novel behavioral setup named INGEsT for investigating the neural dynamics underlying innate ingestive behavior as well as learned operant behaviors. Our data demonstrated that food restricted mice eat pellets at the beginning in the behavioral chamber, and switch between water drinking behavior and feeding behavior after around 15 minutes, indicating physiological body state transition, driven by internal states. In addition, we developed a food or water associated operant behavior task to investigate neural mechanisms of motivation and reward. Furthermore, we developed a novel behavioral task, a self-paced food and water choice task to study neural dynamics underlying different motivations and rewards. Overall, this work develops a behavioral setup benefiting the systems neuroscience field and helping to investigate the neural mechanisms of internal body states, motivation, reward, and choice.

The main aim of developing the INGEsT chamber is to enable the study of self-motivated food/water choice behavior. This is the first published study combining a solid pellet dispenser with a lickometer, allowing us to investigate neural mechanisms of feeding/drinking motivation and food/water reward without any confounds of providing liquid nutrition. Furthermore, the INGEsT chamber can also be used to study very challenging questions, e.g., neural mechanisms of internal body states and memory related to feeding and drinking behaviors.

The INGEsT can precisely record the time point of pellet retrieval and licking water at a time resolution of a microsecond. In the free feeding and drinking behavioral task, a pellet is automatically delivered within a second after the previous pellet retrieval, and water is delivered one drop per second if licking is detected. The amount of water delivered can be adjusted by the open time of the valve. This behavior is therefore ideal for studying slow dynamics since each feeding or drinking bout can last for seconds to a minute.

The INGEsT chamber can also be used for self-paced operant behavior, e.g., pellet or water delivery following an active nose poke. We analyzed the neural activity at different phases of the nose poke and ingestive behavior, i.e., pre-nose poke, pre-pellet retrieval, pre-licking water, nose poke, pellet retrieval, licking water, etc. Here, neurons can be identified as potentially representative of motivation or reward, though additional studies, including activation or inhibition of neural populations would be needed to confirm those correlations. In addition, we also analyzed inactive nose pokes in which no reward is delivered, allowing us to examine whether the neurons only encode nose poke action or not. Our current work did not introduce an interval between nose poke and food/water delivery, which decreased the difficulty of the task. Therefore, in some trials, neural responses to nose pokes cannot be distinguished from a neural response to licking water. However, the chamber allows for flexible training parameters, and in the future, we will introduce a 1 second delay before pellet or water access after a nose poke, which will address the overlap issue. One caveat of this task is that If the mice retrieve the pellet, water will not be available until the next nose poke, but unretrieved pellets can remain in the food well even if mice choose to lick for water. This is because the limited communication ports of the electronic board in the FED3, but in the future, we will improve the hardware to address this.

Reversal learning task is successfully performed in this study. Here, we reverse the active nose poke port, aiming to avoid habitual nose-poke behavior, and did not require mice to perform to a certain reversal criteria. However, the INGEsT chamber can also be used to investigate reversal learning itself, in which case additional sessions would be needed to improve performance.

## Conclusion

This study successfully developed the INGEsT to study neural mechanisms of feeding and drinking behavior in the same experiment. This is a fundamental contribution to the ingestive behavior field and will help researchers to better investigate the underlying mechanisms of basic questions and brain disorders.

## RESOURCE AVAILABILITY

### Lead Contact

Further information and requests should be directed to and will be fulfilled by the Lead Contact (Sarah A. Stern).

### Materials Availability

Materials used here are available from the Lead Contact upon reasonable request.

### Data and Code Availability

Raw data and code supporting the current study are available from the Lead Contact upon reasonable request.

### EXPERIMENTAL MODEL AND SUBJECT DETAILS

#### Mice

All experiments were approved by the Max Planck Florida Institute for Neuroscience Animal Care and Use Committee (Protocol #22-002) and were in accordance with the National Institute of Health Guide for the Care and Use of Laboratory Animals. Maximal efforts were made to reduce the suffering and the number of mice used. Male and female ObRb-Cre (Jackson Laboratory, 008320) and Htr3a-Cre:Ai14 (MGI:5435492, and Jackson Laboratory, 007914) mice are 12–20 weeks old at the beginning of behavioral experiments. Animals (ObRb-Cre) for free feeding and drinking behaviors were kept in individual cages under standard conditions in a day/night cycle of 12/12 hours (lights on at 7 am). For the reversal learning and choice behavior, mice (ObRb-Cre and Htr3a-Cre) were kept in individual cages in a reversed light-dark cycle.

## Materials and methods

### Behavioral setup

| Item | Quantity | Catalog number | Company | Schematic Name | Price (\$) |
| --- | --- | --- | --- | --- | --- |
| **Microcontroller** |  |  |  |  |  |
| Bpod State Machine r1 | 1 | 1027 | Sanworks | Bpod | 925 |
| Bpod State Machine r2 | 1 | 1024 | Sanworks | Bpod | 945 |
| Analog input module 8ch | 1 | 1021 | Sanworks | Analog input module 8ch | 495 |
| Analog output module 4ch | 1 | 1013 | Sanworks | Analog output module 4ch | 495 |
| Bpod Valve Driver Module | 1 | 1015 | Sanworks | Bpod Valve Driver Module | 285 |
| **Lickometer** |  |  |  |  |  |
| Dual Lick Port Detector | 1 | 2021-022 | Janelia | Lick port detector | 80 |
| RJ45 Breakout Board | 2 | ET-CON RJ45 | ETT | RJ45-TERM | 16 |
| Solenoid valve | 2 | LHDA 12332 15H | The Lee Company | Water valve | 50*2=100 |
| Tubing ID 1/16" | 1 | 57217 | Tygon | tubing | 12.5 |
| Tubing ID 1/32" | 1 | 54623 | Tygon | tubing | 17.5 |
| Hypodermic tubing | 1 | 316H18TW | Microgroup | Metal lick port | 45 |
| **Feeding detection** |  |  |  |  |  |
| Fed3 | 1 | Fed3.1 | Open Ephys | Fed3 | 205 |
| **Computer and monitor** |  |  |  |  |  |
| Computer | 1 | Precision 3650 | Dell | Computer | 1109 |
| Monitor | 2 | Dell S2421 | Dell | Monitor | 150*2=300 |
| Monitor mounting kit | 1 | Ergotron kit | Dell | Mounting kit | 134 |
| Camera | 2 |  | ELP | Camera | 60 * 2 = 120 |
| **Optogenetics** |  |  |  |  |  |
| LED | 2 | M470F3 | Thorlabs | LED | 419.27*2=838.54 |
| LED driver | 2 | LEDD1B | Thorlabs | LED driver | 331.64*2=663.28 |
| Power supplier | 2 | KPS201 | Thorlabs | Power supplier | 37.66*2=75.32 |
| Pulse Pal V2 | 1 | 1102 | Sanworks | Pulse generator | 995 |
| **In total:** | | | | | 7856.14\$ |

### Surgery and viral administration

Mice were anesthetized by isoflurane (3% for the induction and 1.5% during the surgery) and placed on a stereotaxic apparatus (Model 900, KOPF instruments, CA, USA) with a mouse adaptor and lateral ear bars. For viral vector delivery, AAV vectors were loaded in a glass pipette and infused by a nanoliter injector (DRUMMOND, nanoinject II). The injection coordinates in anteroposterior (AP) / mediolateral (ML) / dorsoventral (DV) from Bregma, were in mm: for the insular cortex, +0.5/±3.85/-3.9 (250 nL, 100 nL/min), for the ProView Integrated GRIN lens implantation +0.5/±3.85/-3.9 (0.5mm X 6.1mm, 1050-004415). The virus is pGP-AAV-syn-jGCaMP8m-WPRE (Addgene 162375, 1.7x10^13^ GC/mL). The coordinates used were decided according to the mouse brain atlas (Paxinos and Franklin, 2001).

#### Behavioral training

1) Food restriction: Adult mice over 20 grams receive 70% (∼2g) of standard 5V5R chow food on the floor in home cages for around a week and body weight reaches to 85-90% of body weight before animal training.
2) Free feeding and drinking behaviors: free food and water in the behavioral chamber for 3-5 sessions (one session per day, each session lasts 30 minutes); recording calcium signal during the behavior. Mice will be provided extra chow to reach to 70% (∼2g) of daily food if mice eat food pellets (Dustless Precision Pellets® Rodent, Grain-Base #F163) from FED3 feeder less than 70% of daily food during the behavioral task.
3) Nose poke feeding training and reversal learning: Nose poke is required to get available food. After a left/right nose poke followed by an auditory cue, a pellet will be delivered to the pellet well. The response period is 10 seconds. Active nose poke port switches from one side to another side every 5 trials. Active water spout is on the same side as the active nose poke port. Mice can only have available pellets during the behavior. Mice will be provided extra chow to reach 70% (∼2g) of daily food if mice eat less than 70% of daily food during the behavioral task. This training phase will last for 8-10 sessions.
4) Water restriction: Mice receive 0.8 mL water per day and food ad libitum for 7 days before starting animal training.
5) Nose poke drinking training and reversal learning: Nose poke is required to get available water. After a left/right nose poke followed by an auditory cue, 5µl water is available after licking the water spout. The response period is 10 seconds. Active nose poke port switches from one side to another side every 5 trials. Active water spout is on the same side as the active nose poke port. Mice can only have available pellets during the behavior. Mice will be provided extra water to reach 0.8 ml if mice drink less water than 0.8 ml during the behavioral task. This training phase will last for 8-10 sessions.
6) Nose poke training and reversal learning: Nose poke is required to get available food and water. After a left/right nose poke followed by an auditory cue, a pellet will be delivered to the pellet well and 5µl water is available after licking the water spout. The response period is 10 seconds. Active nose poke port switches from one side to another side every 5 trials. Active water spout is on the same side as the active nose poke port. Mice can only have available pellets during the behavior. Mice will be provided extra chow to reach to 70% (∼2g) of daily food if mice eat less than 70% of daily food during the behavioral task. This training phase will last for 8-10 sessions.
6) Food/water choice task: After a nose poke followed by an auditory cue, mice could either get a pellet from pellet well or get 5µl water by licking the active water spout. If mice get pellet first, water will not be available. But if mice lick water first, pellet is still available. This training phase will last for 3∼7 sessions under water restriction conditions, then do the same behavior under food restriction conditions.

## DATA ANALYSIS

Behavioral data was collected by MATLAB. Imaging data was collected by Inscopix DAQ box and preprocessed by Inscopix IDPS software to obtain ΔF/F values. Data analysis was done by MATLAB.

## ACKNOWLEDGEMENTS

We thank the Max Florida Institute ARC, and in particular Colleen Neiner, for animal care. We thank Dr. Carmen Varela and Joseph Wasserman for the discussion of the behavioral task. We also thank Darielle Lewis-Sanders and Elisa Mizrachi for their help in administration. We are grateful to Yingxue Wang and Freda Pang for their help in making and using dual lick port detector. We thank Dr. Hidehiko Inagaki and Shouvik Majumder for their assistance in Matlab programming training. This work was supported by the National Institutes of Health Brain Initiative R00DA04749 (S.A.S) and National Institutes of Health DP2DA060436 (S.A.S) the Brain Research Foundation Seed Grant (S.A.S), the One Mind Foundation Rising Star Award (S.A.S.), the Max Planck Florida Institute for Neuroscience, and the Max Planck Society.

## AUTHOR CONTRIBUTIONS

Z.Z. conceived the project, performed the behavioral and imaging experiments, analyzed behavioral and imaging data, and wrote the original draft. B.X. and Z.Z. wrote codes for data analysis. B.X. analyzed animal video. C.L., S.A., I.M., and A.S. assisted in behavioral experiments. M.B. contributed project supervision regarding Htr3a mice. S.A.S contributed funding acquisition, project supervision, and writing the manuscript. All authors read and approved the manuscript.

